# Wearables-Only Analysis of Muscle and Joint Mechanics: An EMG-Driven Approach

**DOI:** 10.1101/2021.06.16.448524

**Authors:** Reed D. Gurchiek, Nicole Donahue, Niccolo M. Fiorentino, Ryan S. McGinnis

## Abstract

Complex sensor arrays prohibit practical deployment of existing wearables-based algorithms for free-living analysis of muscle and joint mechanics. Machine learning techniques have been proposed as a potential solution, however, they are less interpretable and generalizable when compared to physics-based techniques. Herein, we propose a hybrid method utilizing inertial sensor- and electromyography (EMG)-driven simulation of muscle contraction to characterize knee joint and muscle mechanics during walking gait. Machine learning is used only to map a subset of measured muscle excitations to a full set thereby reducing the number of required sensors. We demonstrate the utility of the approach for estimating net knee flexion moment (KFM) as well as individual muscle moment and work during the stance phase of gait across nine unimpaired subjects. Across all subjects, KFM was estimated with 0.91 %BW•H RMSE and strong correlations (*r* = 0.87) compared to ground truth inverse dynamics analysis. Estimates of individual muscle moments were strongly correlated (*r* = 0.81-0.99) with a reference EMG-driven technique using optical motion capture and a full set of electrodes as were estimates of muscle work (*r =* 0.88-0.99). Implementation of the proposed technique in the current work included instrumenting only three muscles with surface electrodes (lateral and medial gastrocnemius and vastus medialis) and both the thigh and shank segments with inertial sensors. These sensor locations permit instrumentation of a knee brace/sleeve facilitating a practically deployable mechanism for monitoring muscle and joint mechanics with performance comparable to the current state-of-the-art.

## I. Introduction

REMOTE patient monitoring, enabled by advances in wearable technology and algorithms for human movement analysis, promises to improve the assessment and treatment of musculoskeletal disease [1]. Recent work quantifying stride-by-stride gait mechanics at segment-, joint-, and muscle-specific levels has shown that these variables may provide more sensitive measures of patient health than the more typical gross measures of physical activity [2], [3]. Despite these advances, many of the most clinically relevant variables have yet to be observed outside of controlled, laboratory environments. Ideally, assessments would quantify cumulative loading of muscle and articular tissue across every step taken in daily life to best characterize the mechanical stimuli driving tissue adaptation. Characterization at this level could enable personalized therapies and optimal evaluation of intervention efficacy. Further, remote monitoring of these variables could provide novel insight into musculoskeletal disease etiology. For example, in osteoarthritis, load is known to have a positive effect on healthy tissue and yet detrimental effects on diseased tissue [4]. It is not known when this transition occurs, but monitoring cumulative tissue loads under free-living conditions could allow the investigation of these and other cumulative load-dependent phenomena. However, new methods are needed for characterizing joint and muscle mechanics in remote environments to enable these important clinical and research advancements.

The biomechanical variables associated with these analyses (e.g., joint moment, muscle force) provide far more clinical utility than what is typically evaluated remotely (e.g., physical activity, spatiotemporal gait variables). While both frontal and sagittal plane joint moment are important concerning musculoskeletal diseases of the knee, knee flexion moment (KFM) in particular, is thought to play an especially important role in early knee osteoarthritis [5], [6] and for monitoring patients following reconstructive surgery of the anterior cruciate ligament (ACLR) [7]–[10]. It is also critical to characterize individual muscle function. Muscle power, for example, is a well-known determinant of physical function [11] and the phenomena of work- [12] and load- [13] induced muscle hypertrophy motivate tracking cumulative muscle work and force which may provide a basis for optimal exercise prescription and understanding subsequent dose-response relationships [14]. These analyses may be especially relevant for monitoring patients post-ACLR wherein the knee extensor and flexor musculature are compromised [15], [16] due to muscle atrophy [17] and muscle activation deficits [15]. In this case, early intervention is critical [18] and continuous, remote monitoring augments personalized rehabilitation for targeting specific biomechanical outcomes [19].

While physics-based techniques exist for estimating these clinically relevant variables from wearables [20], [21], they require complex sensor arrays that discourage use outside of research contexts [22]. Regression algorithms have been proposed to reduce the number of required sensors [23], but at the expense of generalizability [24]. Further, machine learning techniques do not characterize the dynamics of some relevant internal state variables (e.g., muscle contraction dynamics) which are modeled in physics-based techniques and may be particularly useful in the application of these techniques for rehabilitation. To leverage the strengths of both approaches, hybrid solutions have been proposed wherein machine learning is used only where the physics are not well understood or insufficiently informed [23], [25]. To the authors’ knowledge, only one pilot study [26] has explored a hybrid method wherein KFM was estimated in an electromyography (EMG)-driven simulation. However, validation was for a single subject and machine learning was used to solve for some kinematics that could have been estimated from physics-based techniques (e.g., knee flexion angle from thigh-and shank-worn IMUs [27]).

In the current work, we introduce a new method for characterizing muscle and joint mechanics during walking which utilizes EMG-driven simulation of muscle contraction and inertial measurement unit (IMU)-driven forward kinematics. This approach was designed to enable more effective management of musculoskeletal diseases of the knee joint. We demonstrate the performance of the proposed approach for characterizing KFM as well as individual muscle moment and work by comparison to standard methods.

## II. Proposed Technique

Figure 1 summarizes the proposed technique. The idea is to model each muscle contributing to KFM as a Hill-type actuator and simulate contraction using an EMG-driven approach. The required inputs then are the excitation of each muscle and the length of each muscle-tendon unit (MTU). The novelty of the current work is to implement this approach with a reduced sensor array such that the technique may be feasibly deployed for remote monitoring. To this end, the proposed technique (now referred to as IMC-GP) uses two IMUs (one each on the thigh and shank) to solve for the system kinematics and compute MTU lengths in a process referred to as inertial motion capture (IMC) while the number of required electrodes is reduced by instrumenting only a subset of muscles whereupon the remaining excitation signals (would be unmeasured in practice) are informed by the measured subset using a Gaussian Process (GP) model of the associated muscle synergy functions [28]. In the current implementation, the subset included three muscles: medial (MG) and lateral (LG) gastrocnemius and vastus medialis (VM). These locations were chosen because they are close to the knee joint such that they could instrument a knee brace for practical deployment.

**Fig. 1.**
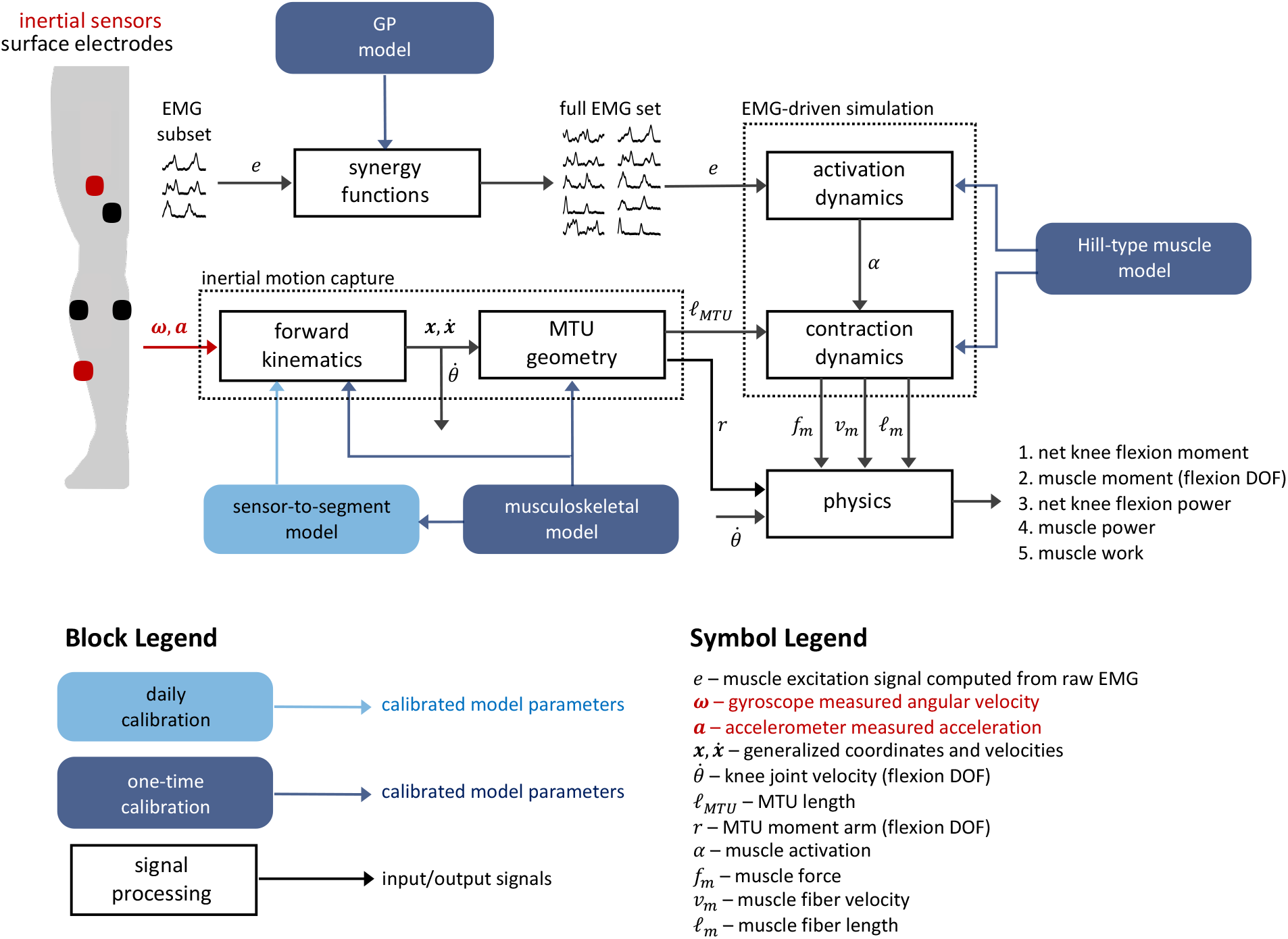
Schematic overview of the proposed technique. Gyroscope and accelerometer data from thigh- and shank-worn IMUs (red sensors, upper left) drive the system kinematics (inertial motion capture) from which the required MTU lengths are computed. EMG data from a subset of instrumented muscles (black sensors, upper left) are mapped to the required full set of excitations through Gaussian Process models of the associated muscle synergy functions. The MTU lengths and excitation signals then drive the simulation of muscle contraction from which the biomechanical variables of interest are computed.

The musculoskeletal model, Hill-type muscle models, and GP models all require a one-time calibration. Calibration of a sensor-to-segment model and EMG normalization is required each time the IMUs and electrodes are attached (e.g., daily). The following sections describe these models and their calibration as well as the IMC analysis, EMG-driven simulation of muscle contraction, and the computation of the biomechanical variables of interest.

### A. Musculoskeletal Model and Calibration

The musculoskeletal model consisted of five segments including a foot, shank, thigh, patella, and pelvis; three joints including a two degree-of-freedom (DOF) ankle, single DOF knee (tibiofemoral), and a three DOF hip; and ten MTUs including the MG, LG, VM, vastus intermedius (VI), vastus lateralis (VL), rectus femoris (RF), long (BFL) and short (BFS) heads of the biceps femoris, semitendinosus (ST), and semimembranosus (SM). MTU path points (origin, insertion, and via points), cylindrical geometry of the femoral condyle, and the origin and insertion of the patellar ligament were taken from Horsman et al. [29]. Average path points were used for multi-element muscles (e.g., superior and inferior elements of the VL). A single via point was used for the MG and LG located at the apex of the shortest curve between origin and insertion points wrapping posteriorly around the cylinder modeling the femoral condyle. The model had 12 mechanical DOFs described by 28 generalized coordinates including translational (n=3) and rotational (n=4, quaternion) coordinates for each segment except the patella (configuration was a function of the knee flexion angle) subject to 16 equality constraints: four enforce the quaternion unit length constraints; nine enforce the non-translating joint constraints; and three enforce the hinge and universal joint constraints on the knee and ankle. The angle of the patella relative to the patellar ligament was modeled as a constant 20° [30]. The angle of the patellar ligament (constant length) relative to the shank was based on the results of van Eijden et al. [30].

Two calibration trials were used to calibrate the subject-specific musculoskeletal model. A static calibration (standing in anatomical position) was used as the reference configuration in which relative positions of segment-fixed points define each rigid body segment including markers (see [31] for marker set) and MTU path points. A functional calibration trial was used to identify joint centers and the knee flexion axis. The subject exercised multiple movements of each joint exciting all joint DOFs through the full range of motion. In this case, segment kinematics were computed independently without regard for any mechanical constraints: orientation via Davenport’s solution to Wahba’s problem [32] wherein every unique two-marker combination for all segment-fixed markers during the static calibration trial were used as the reference vectors (weighted by their length squared) and position as done by Spoor and Veldpaus [33]. Hip and ankle joint centers were estimated using the pivoting algorithm [34] and the knee flexion axis using the SARA method [35]. Knee joint center was defined as the point on the knee flexion axis closest to the femoral epicondyle midpoint.

Segment-fixed coordinate systems were constructed with basis vectors coincident with the principal axes of inertia and origin with the segment center of mass (inertial parameters taken from [36]). Local MTU path points were scaled based on an anthropometric measure of each segment relative to the same from the data reported in [29] (e.g., segment length). Knee extensor and medial hamstring insertion points were adjusted to better align their knee flexion moment arms with published data [37], [38].

### B. Sensor-to-Segment Model Calibration

The daily sensor-to-segment model calibration requires three calibration trials: the same static and functional (hip/knee joint) calibration trials used for calibrating the musculoskeletal model and straight walking. The system configuration during the daily static calibration is assumed equivalent to the reference configuration. The TRIAD algorithm [39] was used to determine the orientation of the IMU frames relative to the segment frames. Reference vectors were the knee joint flexion axis and gravity vector with full trust given to the former. The representation of these vectors in the segment frames were taken from the reference configuration. The representation of the gravity vector in each IMU frame was computed as the average direction of the accelerometer output during the static calibration trial. The representation of the knee joint flexion axis in each IMU frame and the position of the knee joint center relative to each IMU were determined using a nonlinear least-squares method [27], [40]. Data recorded during the hip and knee joint movements of the daily functional calibration trial were used for both calibrations (knee joint flexion axis and knee joint center) in addition to the walking calibration trials for calibrating the knee joint flexion axis.

### C. Inertial Motion Capture

Shank and thigh IMU data were first expressed relative to their segment coordinate systems based on the calibrated sensor-to-segment model. The medio-lateral component of the shank gyroscope signal was used to identify foot contact and foot off events [41]. Shank accelerometer data were used to identify the most still quarter of the identified stance phase (interval for which the sum of the accelerometer signal variances was least) during which the average acceleration was used to estimate shank orientation at the middle of the interval (assuming zero heading angle) [42]. Shank orientation before and after this mid-stance instant was obtained using the analytic solution to the quaternion kinematic equation [43] following an assumption of constant angular rate between measured samples equal to the average of the two samples. Knee flexion angle (*θ*) was estimated using an RTS Kalman smoother [44] implementation of a complementary filter [27]. Thigh orientation was determined from shank orientation and knee flexion angle. Pelvis orientation was assumed neutral except that the heading angle was constant and equal to the average shank heading angle during stance. The acceleration of the knee and ankle joint centers was computed from shank accelerometer data (after removing gravity) along with shank gyroscope-measured angular velocity and the known joint center position relative to the shank IMU [45]. Ankle joint center position was computed by double (trapezoidal) integration of ankle joint center acceleration. Foot heading angle was equivalent to that of the shank, roll angle was zero, and pitch angle was computed based on a simple foot-ground contact model (Fig. 2). Given all segment orientations and the ankle joint center position throughout stance phase, the remaining generalized coordinates were given from the calibrated musculoskeletal model. MTU length and knee flexion moment arm were computed as in [46].

**Fig. 2.**
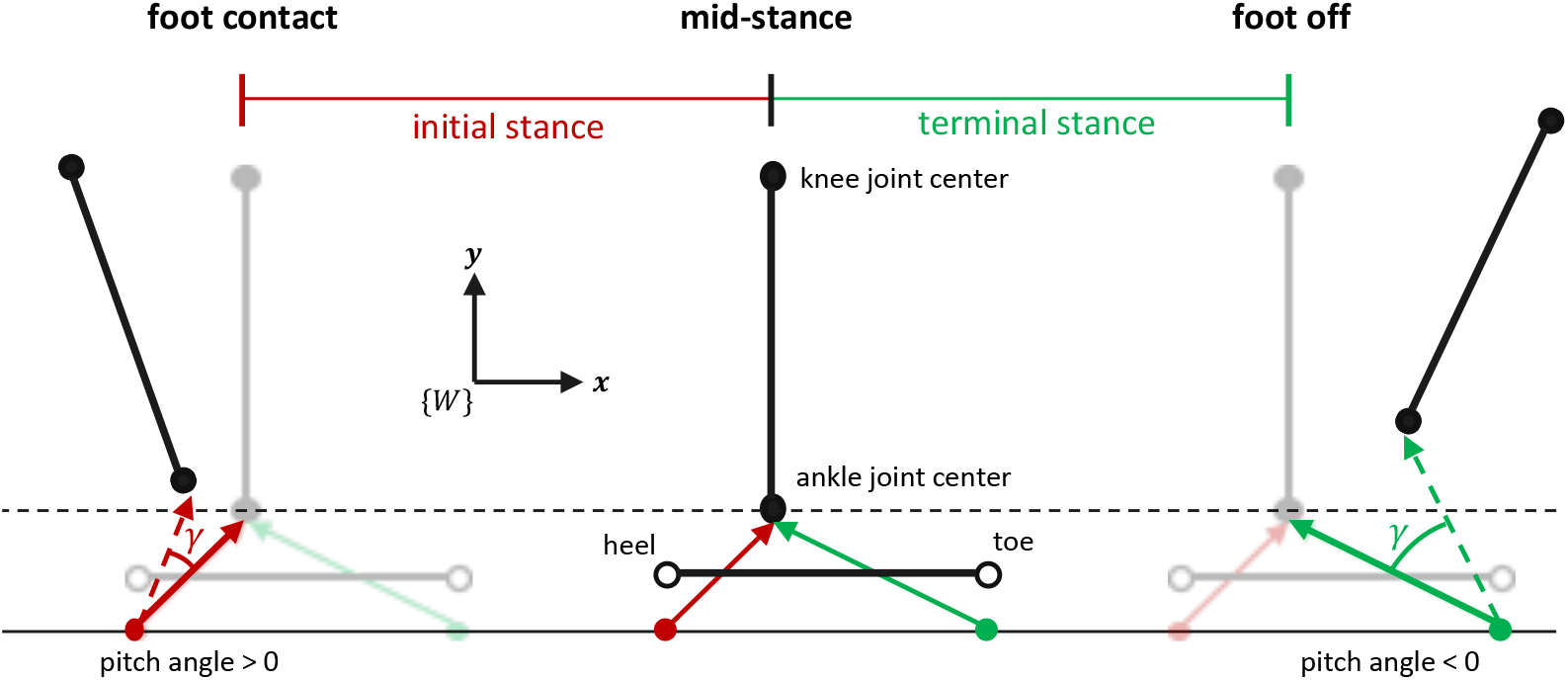
Foot-ground contact model for estimating foot pitch angle (*γ*) in the IMC analysis. Ankle joint center position during stance was available as described in the main text. Planar translation was assumed and thus only the world frame {*W*} vertical and horizontal coordinates were needed. Ankle joint center height at mid-stance (middle) was set equal to that in the reference configuration (black dashed line) from which the positions of the initial stance rotation point (red dot, inferior to the heel) and terminal stance rotation point (green dot, inferior to the toe) were computed. Assuming rotation was about these points in the respective intervals, *γ* was computed as the angle between the rotation point-to-joint center vectors at each time instant during initial stance (red dashed arrow) or terminal stance (green dashed arrow) and the same in the reference configuration (solid red and green arrows at mid-stance).

### D. EMG-Driven Simulation of Muscle Contraction

MTU geometry was modeled as in [47] such that pennation angle (*ϕ*) and tendon length (ℓ_*T*_) were explicit functions of MTU(ℓ_*MTU*_) and muscle fiber (ℓ_*m*_) length as per

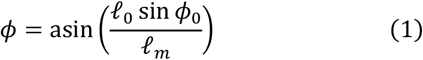

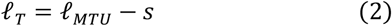

where *s* (equal to the product ℓ_*m*_ cos *ϕ*) is the projection of the fiber length onto the MTU, ℓ_0_ is the optimal fiber length, and *ϕ*_0_ is the pennation angle when ℓ_*m*_ = ℓ_0_ (taken from [29]). Tendon force (*f*_*T*_) was modeled similar to [48] as per

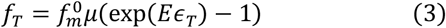

where 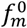 is the maximal isometric force of the muscle, *μ* is constant and equal to exp(−0.04*E*), *ϵ*_*T*_ is the tendon strain modeled as a function of ℓ_*T*_ and tendon slack length (ℓ_*s*_) [49], and the parameter *E*(*df*_*T*_/*dϵ*_*T*_ when *ϵ*_*T*_ was set to 35.00. Muscle force projected onto the *f*_*T*_ line of action (*f*_*m*_) was modeled as per [47]

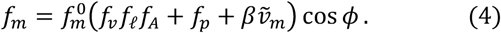

The parenthesized term in (1) is the normalized muscle force which is scaled by 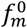 and projected onto the *f*_*T*_ line of action via multiplication with cos *ϕ*. The functions *f*_*v*_, *f*_ℓ_, and *f*_*p*_ model the force-velocity, active force-length, and passive force-length properties of muscle, respectively. Fiber length ℓ_*m*_ normalized by ℓ_0_ is input to both *f*_ℓ_ and *f*_*p*_. The input to *f*_*v*_ is fiber velocity (*v*_*m*_) normalized by the maximal fiber shortening velocity (set to 15 optimal fiber lengths per second [50]); denoted 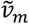. In this study, *f*_*v*_ and *f*_ℓ_ were both modeled as in [48] and *f*_*p*_ as in [51] with passive muscle strain due to maximal isometric force (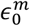 in [51]) set to 0.55. The input to the activation nonlinearity function (*f*_*A*_)), modeling the nonlinear relationship between activation and muscle force [52], [53], is the activation signal (*α*), the dynamics 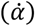 of which were driven by the muscle excitation signal (*e*). The output of *f*_*A*_ is also used in *f*_ℓ_ to model the dependency of the optimal fiber length on muscle activation as in [54]. The product 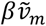 models damping effects within the fiber where the coefficient (*β*) was set to 0.01 [55]. Several parameters not yet specified (e.g., 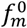, *f*_*A*_, 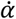, ℓ_0_, ℓ_*s*_) were determined in a calibration process described in section II.F.

In the current implementation, fiber length ℓ_*m*_ and muscle activation *α* were the state variables for the muscle contraction dynamics and system inputs were MTU length ℓ_*MTU*_ and muscle excitation *e* (computed from raw EMG as in [54]). The excitation of the BFS, SM, and VI was assumed equivalent to that of the BFL, ST, and the average of VM and VL, respectively. The equivalence between the tendon (*f*_*T*_) and muscle (*f*_*m*_) force gives rise to the equilibrium equation

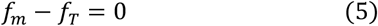

which is an implicit formulation for the dynamics of the fiber length state variable [47], [54], [56]. An implicit solver (ode15i in Matlab) was used to numerically integrate (5). Initial conditions for ℓ_*m*_ and *v*_*m*_ consistent with (5) were found numerically (decic in Matlab). This required an initial guess that may not satisfy (5) for which a rigid tendon was assumed. Activation dynamics were simulated using a Runge-Kutta formula (ode45 in Matlab).

### E. Computation of the Biomechanical Variables of Interest

Net KFM was computed as the sum of the flexion moments generated by each MTU given by the product of *f*_*m*_ and the knee flexion moment arm. Cumulative concentric (*W*_*con*_) and eccentric (*W*_*ecc*_) work were computed using a trapezoidal approximation to the line integral

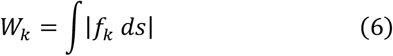

for *k* ∈ {*ecc, con*} where |·| denotes absolute value and

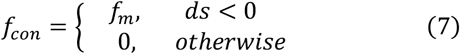

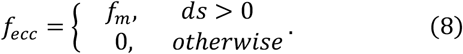

### F. Gaussian Process Model Calibration

The proposed technique uses excitations computed from the raw EMG of the three instrumented muscles to estimate the other four (unmeasured in practice). To this end, four GP models were trained (using open-source toolboxes [28], [57]) to approximate the four associated muscle synergy functions [28]. Input muscles were VM, MG, and LG. Input window length was 1.50 s with a 0.75 s window relative output time. The GP model covariance function was the isotropic squared exponential and the mean function was constant [28].

### G. Hill-Type Muscle Model Calibration

A set of calibration walking trials from which a ground truth estimate of KFM based on inverse dynamics (ID) and a reference EMG-driven (referred to as OMC-Full) estimate were required for identifying Hill model parameters. Biomechanical variables were computed via OMC-Full in the same way as for IMC-GP except kinematics were solved using optical motion capture (OMC, described below) and a full set of measured excitations as opposed to the three-muscle subset.

#### 1) Inverse Dynamics

The 28 generalized coordinates were found by minimizing a squared-error objective [58] (errors between model-based and measured marker positions) subject to the 16 mechanical constraints described previously. The optimal solution was found using the interior-point algorithm (fmincon in Matlab) with analytic Jacobian and Hessian matrices of the objective and constraint equations. The constraint tolerance was set to 1e-6 and all markers were weighted equally. Segment linear and angular velocities and accelerations were approximated using the 5-point central difference method (quaternion velocities were computed first and mapped to angular velocities [43]) and were low-pass filtered using a 4^th^ order, zero-phase, Butterworth filter (6 Hz cutoff frequency) with double-pass adjustments [59]. The knee flexion moment arm and length of each MTU were computed as for the IMC analysis. Intersegmental forces and moments were computed using the recursive Newton-Euler algorithm and KFM by projecting knee intersegmental moment onto the flexion axis.

#### 2) Parameter Optimization

Several physiological parameters related to the contraction dynamics must be optimized for each person and muscle (usually via global optimization [49]). A novelty of the current work is the inclusion of categorical parameters in the tunable parameter set. Specifically, we use Bayesian optimization to optimize two functions, the activation nonlinearity function *f*_*A*_ and the model of the activation dynamics 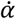, in addition to five scalar parameters. Optimal fiber length ℓ_0_ and tendon slack length ℓ_*s*_ are often included in the tunable parameter set. However, to prevent overfitting [49] we chose to reduce the number of tunable parameters (which would otherwise be larger by inclusion of *f*_*A*_ and 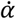) by removing ℓ_0_ and ℓ_*s*_. Instead, ℓ_0_ and ℓ_*s*_ were optimized similar to previous work [60] so that the range of ℓ_*m*_ normalized by ℓ_0_ during walking gait would be within the range 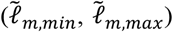 of published data [50], [61] if a rigid tendon model were used. In this case,

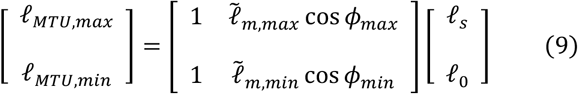

where 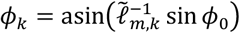 for *k* ∈ {*max, min*} and the range of MTU lengths (ℓ_*MTU,min*_, ℓ_*MTU, max*_) were subject-specific and taken from the walking calibration trials. The solution to (9) yields the optimal ℓ_0_ and ℓ_*s*_.

Bayesian optimization was used to tune a scalar that scaled the maximal isometric force 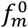 and five muscle activation parameters: activation dynamics model 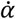, activation time constant *τ*_*a*_, activation-to-deactivation ratio *τ*_*a*_/*τ*_*d*_ where *τ*_*d*_ is the deactivation time constant, activation nonlinearity function *f*_*A*_, and a parameter in *f*_*A*_. Muscles were grouped such that those belonging to the same group were assumed to have the same properties. Similar to previous work [54], three groups were permitted based on MTU structure and function: knee extensors, hamstrings, and gastrocnemii. Further, due to the association between fiber type distribution and the activation-force relationship [53], we required muscles in the same group to have a similar proportion of type I (slow-oxidative) fibers [62]. All muscles within the three groups described previously with this fiber type proportion less than 60% were placed in a new group as were those greater than or equal to 60%. This was the case only for the hamstrings where SM and ST were both 50% type I, while BFL and BFS were both 65%. Thus, the four muscle groups were the knee extensors (VL, VM, VI, RF), lateral (BFL, BFS) and medial (SM, ST) hamstrings, and gastrocnemii (MG, LG) yielding 24 total tunable parameters.

The strength scalar (range: 0.5 – 2.0) scaled 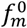 initialized as the product of the physiological cross-sectional area [29] and the muscle stress when 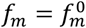 (set to 0.30 MPa). Five activation dynamics models were considered (see appendix for details): (1) a 1^st^ order, linear model [63], (2) a 1^st^ order, nonlinear, piecewise-continuous model [48], (3) a 1^st^ order, bilinear model [64], (4) a 2^nd^ order, linear model [65], and (5) a piecewise version of model (4). All models were unity gain with an electromechanical time delay (40 ms [54]). The range of *τ*_*a*_ was 10-60 ms and of the ratio *τ*_*a*_/*τ*_*d*_ was 0.25-1.00. Three functions were considered for *f*_*A*_ (see appendix for details): (1) an exponential model [54], (2) the A-model [66], and (3) the twice differentiable A-model [67].

The objective function was the average normalized mean squared error between the ID and OMC-Full estimate of KFM across all calibration walking trials where normalization was by the variance of the ID estimate. Optimization was executed using bayesopt in Matlab with the expected-improvement-plus acquisition function, 0.5 exploration ratio, 96 seed points (four times the number of parameters), 300 GP active set size, and the number of maximum objective function evaluations was set to 576 (the number of parameters squared).

## III. Experimental Validation

### A. Data Collection

The proposed technique was validated on nine unimpaired subjects (four female, age: 21 ± 1 years, height: 1.77 ± 0.11 m, mass: 72.10 ± 12.30 kg). Each subject performed 10 overground walking trials at a self-selected normal speed (1.36 ± 0.20 m/s) for which the right foot completely contacted the force plate for a single contact. Thus, one stance phase for the right leg was analyzed per trial. Marker position data were captured using 19 cameras (Vicon Motion Systems, Oxford, UK, 100 Hz). Force plate (AMTI, Watertown, MA, USA) and raw EMG (BioStamp, MC10, Inc., Cambridge, MA, USA) data were collected at 1000 Hz. Electrodes were placed over the MG, LG, VM, VL, RF, ST, and BFL according to SENIAM recommendations [68]. Force plate data were downsampled to 100 Hz for synchronization with marker data as were EMG data after excitations were computed. Shank and thigh IMUs (BioStamp, MC10, Inc., Cambridge, MA, USA, gyroscope range: ± 2000°/s, accelerometer range: ± 16g, 250 Hz) were placed over the distal lateral shank and anterior thigh, respectively. IMU data were downsampled to 100 Hz and digitally time synchronized with marker position data. All subjects provided written consent to participate and all activities were approved by the University of Vermont Institutional Review Board (#18-0518).

Each subject performed the static and functional calibration trials described previously for calibrating the musculoskeletal model. The first seven overground walking trials were set apart for calibrating the MTU parameters and GP models and the last three (test trials) were set apart for validation. The sensor-to-segment model calibration trials were the same static and functional calibration trials used for calibrating the musculoskeletal model and the test trials were used as the walking calibration trials. This mimics how calibration would be done in practice: the identified walking activity being evaluated would also be available for calibration.

### B. Statistical Analysis

Statistics characterizing performance of IMC-GP are reported only for the three test trials. Net KFM from IMC-GP was compared with both ID and OMC-Full and individual muscle moment was compared between IMC-GP and OMC-Full using Pearson’s correlation coefficient (*r*) and root mean square error (RMSE). RMSE was expressed as a percentage of the average range of the two time-series being compared (denoted %range) and of the product of subject body weight (in N) and height (in m); denoted %BW•H. To compare our results to previous work [20], [21], *r* and RMSE were computed for every time-series and then averaged. Average correlations were corrected using Fisher’s z transformation [69] and interpreted as weak (*r* ≤ 0.35), moderate (0.35 < *r* ≤ 0.67), strong (0.67 < *r* ≤ 0.90), and excellent (*r* > 0.90) [21]. Peak knee extension moment (KEM) during initial stance was compared between IMC-GP and ID using *r*, mean absolute error (MAE), and Bland-Altman analysis: mean error (ME) and 95% limits of agreement (LOA) with compensation for repeated measures [70]. Pearson’s *r* was used to evaluate the sensitivity of the IMC-GP analysis to variation in muscle work across subjects by comparison to the OMC-Full analysis. Work was considered only for the contraction type (eccentric or concentric) in which each muscle did the most work.

## IV. Results

All three techniques yielded the same general trend in the KFM time-series (Fig. 3). This was supported statistically by strong to excellent correlations between IMC-GP estimates with those from ID (range: *r* = 0.68-0.96, average: *r* = 0.87) and OMC-Full (range: *r* = 0.74-1.00, average: *r* = 0.95) with RMSE less than 1.00 %BW•H (Table I). Correlations between IMC-GP and OMC-Full estimates of individual muscle moment (see online supplementary material for a graphical comparison) were strong to excellent (*r* = 0.81-0.99) across all muscles with RMSE between 6.46-26.33 %range (Table I). Peak KEM was estimated to within 0.57 %BW•H MAE of the ID estimate (ME: −0.22 %BW•H, LOA: −1.54 to 1.11 %BW•H) with excellent (*r* = 0.92) correlation (Fig. 4). Excellent correlations were also observed between IMC-GP and OMC-Full estimates of cumulative muscle work across all muscles (Fig. 5) except for the VL (*r* = 0.88).

**Fig. 3.**
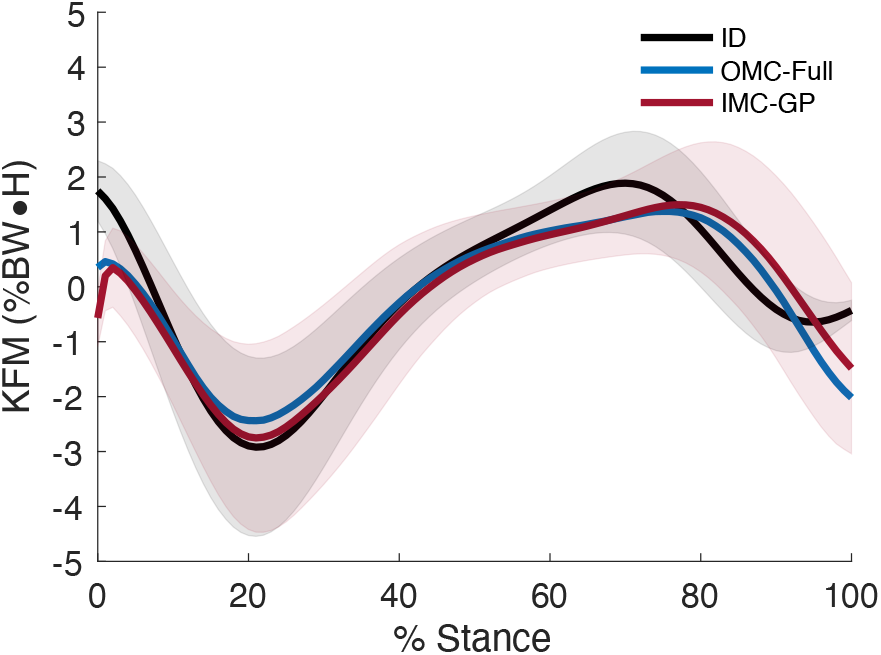
Ensemble average net KFM. Positive (negative) values indicate a flexion (extension) moment. The shaded area is ± 1 s.d. (ID, IMC-GP).

**Fig. 4.**
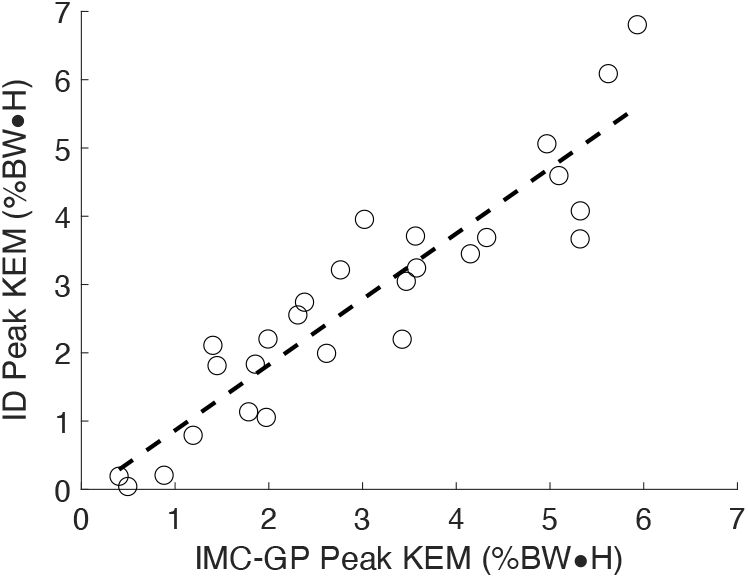
Scatter plot of peak KEM from the proposed technique (IMC-GP) compared to the ground truth inverse dynamics (ID) analysis.

**Fig. 5.**
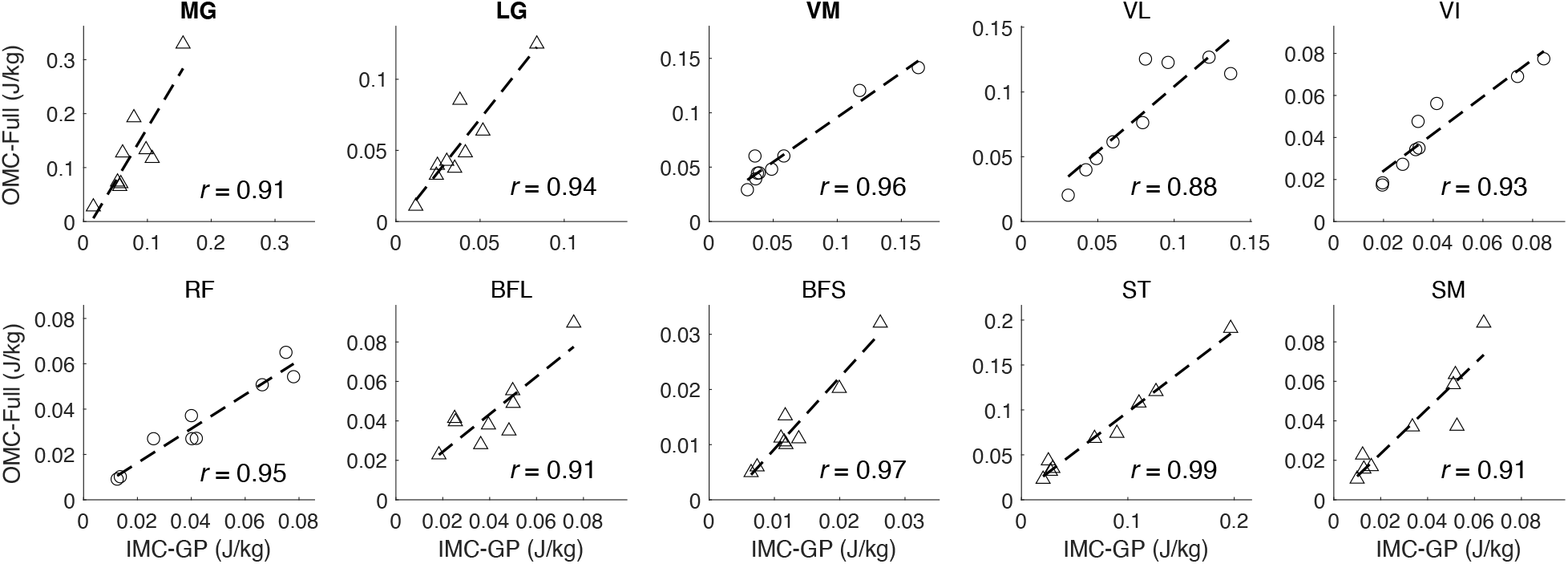
Scatter plots of cumulative muscle work (three-trial average) from the proposed technique (IMC-GP) compared to the reference EMG-driven technique (OMC-Full). Work is shown for the type of contraction in which each muscle did the most work: concentric (triangles) or eccentric (circles). Bold titles indicate muscles for which measured excitations were used to simulate contraction whereas the others were based on the GP synergy function models.

## V. Discussion

The most promising result from this validation was the estimation of KFM with strong correlations (*r* = 0.87) and low errors (0.91 %BW•H, 18.25 %range RMSE) using IMC-GP. These results compare favorably with the current state-of-the-art in physics-based, wearables-only techniques. For example, Karatsidis et al. present an IMU-driven inverse dynamics analysis (17 IMUs, 17 segments) and optimization-based muscle force estimation [20]. KFM was estimated for 11 healthy men across three walking speeds with a moderate correlation (*r* = 0.58) and 1.9 %BW•H (29.8 %range) RMSE. Dorschky et al. present an approach based on optimal control of a musculoskeletal model wherein state variables tracked measured sensor signals (seven IMUs, seven segments) via trajectory optimization [21]. KFM (full gait cycle) was estimated for 10 healthy men across three walking speeds with strong correlations (*r* = 0.81) and 1.5 %BW•H (27.1 %range) RMSE. Compared to these methods, the proposed technique presents a significant reduction in sensor array complexity with comparable estimation performance.

**TABLE I.**
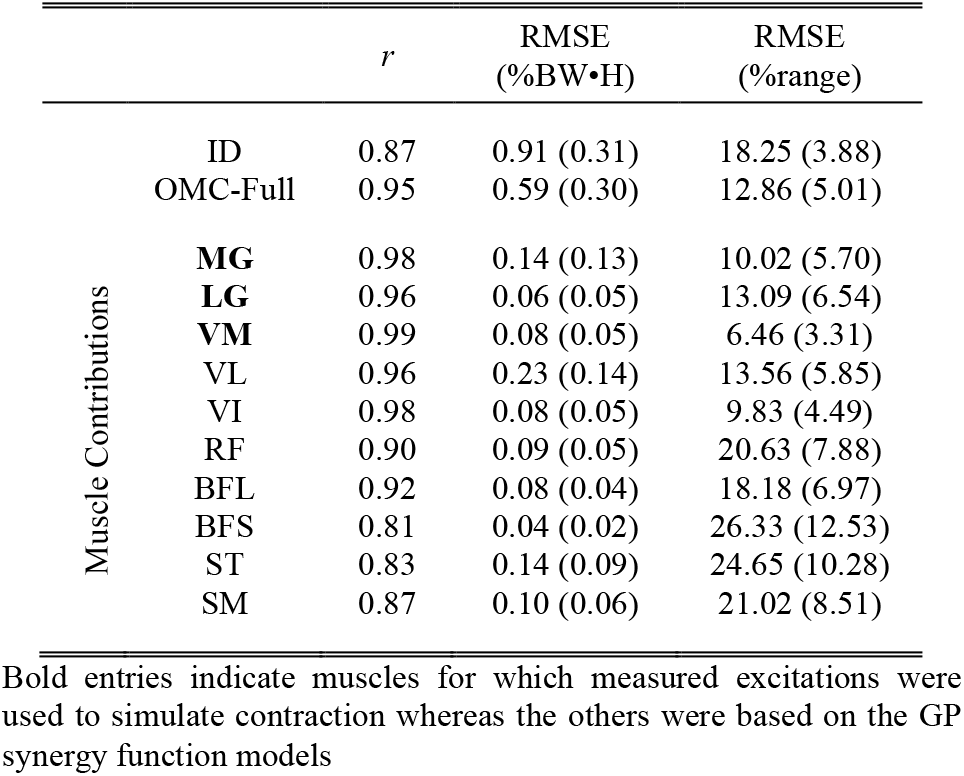
Estimation of Net KFM and Individual Muscle Moment

The proposed technique also compares well with machine learning techniques. For example, estimation of KFM from the proposed technique was comparable to a neural network (NN) using EMG inputs (*r*^2^ = 0.81 vs. 0.76 for IMC-GP) [71] and a linear model using data from an instrumented insole (*r* = 0.89) [72]. In more recent developments, NN-based architectures with IMU inputs have been used to estimate KFM with 1.14 %BW•H RMSE and *r* = 0.98 in a four-sensor, four-segment configuration [73] and with 18.4 %range RMSE and *r* = 0.72 in a two-sensor, two-segment configuration [74]. In addition to characterization of KFM, IMC-GP provides complementary insight into the function and loading of individual muscles which are not modeled in machine learning techniques.

One study (a recent conference paper [26]) has presented results for a hybrid approach similar to IMC-GP: machine learning informed both MTU kinematics and the mapping from an EMG subset to a full set. KFM (full gait cycle) was estimated with 26, 30, and 26 %range RMSE for walking at 1.5, 3.0, and 5.0 km/h, respectively, compared to 18.25 %range RMSE in the current study.

Peak KEM during initial stance was estimated to within 0.57 %BW•H MAE (i.e., 9.98 %BM, percentage body mass) which is less than observed inter-limb differences for patients post-ACLR (1.08 %BW•H [7]), differences between patients post-ACLR and healthy controls (17 %BM [8], 1.72 %BW•H [7], 1.10 %BW•H for patellar tendon graft [9]), and gender differences observed for patients 12 months post-ACLR (33.50 %BM [10]). Therefore, the proposed technique appears able to detect clinically meaningful differences. From at least one study [8], the observed LOA and 9.98 %BM MAE may appear too large to observe differences pre- and post-ACLR. Nevertheless, the observed excellent correlation (*r* = 0.92, Fig. 4) suggests the proposed technique is sensitive to variation in peak KEM and thus could track patient recovery post-ACLR.

The proposed technique is dependent on EMG-driven simulation of muscle contraction and thus the OMC-Full analysis represents the theoretical ceiling of performance. Our comparison of IMC-GP to OMC-Full supports the proposed technique as a promising surrogate for EMG-driven analyses in remote monitoring applications (Table I). Estimation was best for the instrumented muscles (VM, MG, and LG in the current implementation). In a post-hoc comparison, we found that accurate estimation of individual muscle moment was due in large part to accurate estimation of muscle activation (from the GP models) based on similar RMSE (%range) of the two signals: 13.41 in activation RMSE vs. 13.56 in KFM RMSE for VL, 17.25 vs. 20.63 for RF, 25.63 vs. 24.65 for ST, and 25.81 vs. 18.18 for BFL. For instrumented muscles, estimation was better for VM than for MG and LG (Table I). IMU-driven forward kinematics likely explain this discrepancy wherein estimation of knee flexion angle (*r* = 0.98, 4.08° RMSE) was better than for ankle dorsiflexion (*r* = 0.53, 9.93° RMSE). Knee flexion angle was estimated using a Kalman smoother implementation of a previously validated complementary filter [27]. However, to avoid the need for a foot-worn sensor, ankle dorsiflexion was given following a simple foot-ground contact model (Fig. 2). A more complex contact model (e.g., including toes) and/or a data fusion approach (e.g., fusing contact model estimates with a forward dynamic estimate driven by ankle muscles) may improve estimation. Still, the strong correlations and relatively low errors motivate use of IMC-GP for evaluating relative muscle contributions to KFM which has clinical implications for managing musculoskeletal disease.

The results of the correlation analysis suggest the proposed technique was sensitive to variation in muscle work from the reference EMG-driven analysis (Fig. 5). Muscle work is a known stimulus for hypertrophy [12] and objectively quantifies exercise intensity. Thus, the proposed technique points toward continuous monitoring of individual muscle loading in daily life. This could be transformative for personalized therapy enabling novel patient profiling (e.g., characterizing patient-specific exercise dose-response relationships over time), evaluation of intervention efficacy, and the potential to adapt loading prescriptions for managing tissue over- and under-loading. While we demonstrated the characterization of individual muscle function via quantification of moment and work, other variables are necessarily characterized (see online supplementary material) including joint, MTU, and muscle kinematics, power, and force. This thorough characterization of the musculoskeletal system is due to the physics-based nature of the proposed approach and is nonexistent in machine learning alternatives. For the latter, all desired outcome variables are modeled separately and physical relationships between inputs and outputs are not necessarily maintained.

Several limitations should be considered. Validation was only for unimpaired subjects walking at self-selected normal walking speeds. However, walking speeds ranged from 0.88-1.76 m/s which is similar to the range of speeds used in previous multi-speed validations [21] and the range of stride times (0.98-1.13 s) encompass the majority of those observed in free-living, unimpaired gait [75]. Future work should investigate performance across multiple conditions (e.g., multi-speed, multi-load) and in impaired populations. EMG-driven simulation of muscle contraction requires normalized excitations (by maximal voluntary contraction) which may vary throughout the day due to changes in the properties of the skin-electrode interface [76]. Thus, compensatory methods must be developed that are robust to these variations. The knee was modeled as a single DOF hinge joint. However, more complex models with translational DOFs may be more appropriate [77]. Only flexion moments were estimated and muscles with negligible contribution were ignored as in previous studies [78]. Modeling additional musculature would enable analysis of other DOFs and joint contact forces. The foot-ground contact and GP muscle synergy function models assumed walking gait. This would not be problematic for remote monitoring as walking activity is identified in the processing pipeline [2]. However, analysis of other tasks would require development of other task-specific models.

## VI. Conclusion

This study presents a hybrid machine learning- and physics-based technique for analysis of muscle and joint mechanics during walking using only wearable sensor data. Machine learning was used to reduce the number of required surface electrodes for EMG-driven simulation while data from two IMUs drove the system kinematics via physics-based techniques. The proposed technique performed well compared to laboratory standard inverse dynamics and EMG-driven analyses with comparable performance to other wearables-only techniques with more complex sensor arrays. The proposed approach may be easily generalized for analysis of other muscles and joints not considered herein. Importantly, the proposed technique allows sensor placement near the knee joint such that they could be integrated into a knee brace or sleeve for practical deployment. Future work should target remaining barriers to deployment including EMG normalization for long-duration recordings and integrating the required sensors into a wearable accessory.

## Supporting information

Appendix A: Function Sets

Appendix B: Supplementary Analysis

## Acknowledgments

RSM reports stock ownership in Epicore Biosystems, Inc.; Impellia, Inc.; and Allostatech, LLC. RSM reports research funding from MC10, Inc.; Epicore Biosystems, Inc.; US National Science Foundation; US National Institute of Health.

## Notes

This work was supported in part by the Vermont Space Grant Consortium through the NASA Cooperative Agreement under Grant NNX15AP86H.

