## Appendix A: Function Sets for "Wearables-Only Analysis of Muscle and Joint Mechanics: An EMG-Driven Approach"

### Appendix A: Activation Dynamics and Activation Nonlinearity Functions

This study differs from previous work in the use of Bayesian optimization for tuning MTU parameters. Several activation dynamics [1]–[4] and activation nonlinearity functions [5]–[7] have been proposed in previous research and it was not clear which model was most appropriate. The approach in previous work has been to choose a model for the activation dynamics and activation-force nonlinearity and assign it to every muscle. The parameters of those models are then optimized via global optimization (e.g., simulated annealing [5], [8]), often in addition to the optimal fiber length and tendon slack length. However, empirical evidence suggests activation dynamics may be muscle specific and related to fiber type distribution [9], which informed the activation muscle groupings in the current study. Moreover, the activation dynamics have been recognized as the least well established in the Hill model [10]. Further uncertainty was introduced surrounding the activation dynamics by using GP synergy functions to estimate some of the excitation signals. Thus, we chose to include these functions as tunable (categorical) parameters in the global optimization for which Bayesian optimization is suitable.

Five activation dynamics models were considered. The first was a 1st order, linear model ($L_1T_1$) based on that used by Winters and Stark [2],

$$\dot{\alpha} = \bar{\tau}_a^{-1}(e - \alpha) \tag{A.1}$$

where $\bar{\tau}_a$ is the average of the activation and deactivation time constants. The second was a 1st order, nonlinear, piecewise-continuous model ($N_1T_2$) based on that used by De Groote et al. [1],

$$\dot{\alpha} = \frac{1}{2}\left[\frac{(1 + \tanh[0.1(e-\alpha)])}{\tau_a(0.5 + 1.5\alpha)} + \frac{(0.5 + 1.5\alpha)(1 - \tanh[0.1(e-\alpha)])}{\tau_d}\right](e - \alpha). \tag{A.2}$$

The third was a 1st order, bilinear model ($B_1T_2$) based on that used by He et al. [3],

$$\dot{\alpha} = [(\tau_a^{-1} - \tau_d^{-1})e + \tau_d^{-1}](e - \alpha). \tag{A.3}$$

The fourth was a 2nd order, linear model ($L_2T_1$) based on the findings of Milner-Brown et al. [4],

$$\ddot{\alpha} = -2\Omega\dot{\alpha} - \Omega^2\alpha + \Omega^2 e \tag{A.4}$$

where the natural frequency $\Omega$ is equal to the average value of $\tau_a^{-1}$ and $\tau_d^{-1}$. The fifth was a piecewise version of the model (A.4) ($L_2T_2$) where $\Omega = \tau_a^{-1}$ for $\alpha \geq 0$ (during activation) and $\Omega = \tau_d^{-1}$ for $\dot{\alpha} < 0$ (during deactivation). Three activation nonlinearity functions were considered. The first was the exponential model used by Lloyd and Besier [5] (Aexp) based on the findings of Potvin et al. [11],

$$f_A = \frac{\exp(A\alpha) - 1}{\exp(A) - 1}. \tag{A.5}$$

The second was the piecewise A-model developed by Manal and Buchanan [7] (A), based on the findings of Woods and Bigland-Ritchie [9] which is dependent on a single parameter $A'$. The third was a modified version of the (twice differentiable) approximation to the piecewise A-model used by Meyer et al. [6] (Ac),

$$f_A = (1 - A'')\alpha + A''\left(1 + \frac{b_1}{b_2 + b_3(\alpha + b_4)^{b_5}}\right) \tag{A.6}$$

where $b_1 = -7.623$, $b_2 = 4.108$ $b_3 = 29.280$, $b_4 = 0.884$, $b_5 = 17.227$. The parameter $A''$ in (A.6) was a function of $A$ such that the derivative of (A.6) with respect to $\alpha$ evaluated at $\alpha = 0$ was equivalent to that of (A.5). A similar adjustment was made to the single parameter $A'$ of the piecewise A-model such that $A' = \frac{0.12}{0.35}A''$. These parameter adjustments were made so that the single tunable parameter $A$ yielded similar behavior in all three activation nonlinearity functions for the range $-3 \leq A < 0$ [5]. The step response for each activation dynamics model and the behavior of the three activation nonlinearity functions is shown in Fig. A.1.

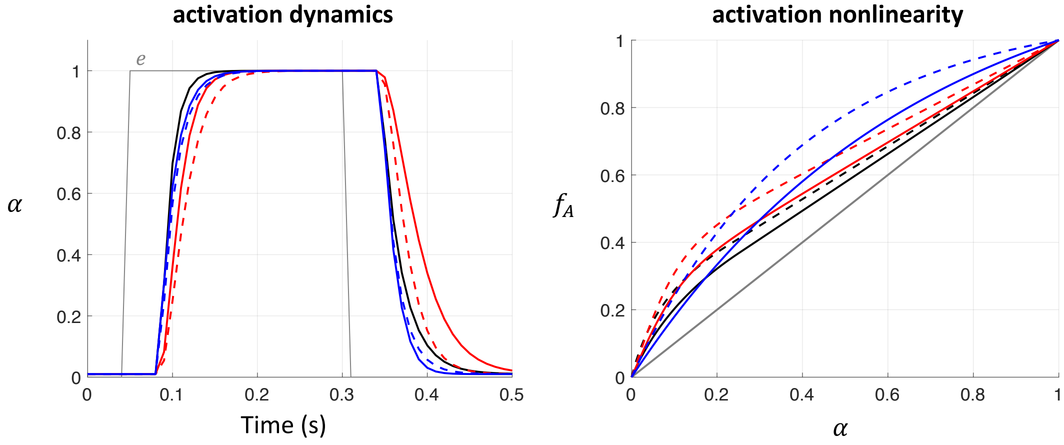


Fig. A.1. Activation dynamics and activation nonlinearity functions included in the tunable parameter set of the Bayesian optimization. The step response for each activation dynamics model is shown in the left plot. The electromechanical time delay was set to 40 ms, $\tau_a$ was set to 12 ms and the ratio $\tau_a/\tau_d$ was set to 0.5 such that $\tau_d$ = 24 ms. The solid black line is the $B_1T_2$ model, the solid blue line is the $N_1T_2$ model, the dashed blue line is the $L_1T_1$ model, the solid red line is the $L_2T_2$ model, and the dashed red line is the $L_2T_1$ model. The behavior of the activation nonlinearity functions is shown on the right where solid lines correspond to $A$ = -1.50 and dashed lines to $A$ = -2.50. The black lines correspond to the A model, the red lines correspond to the Ac model, the blue lines correspond to the Aexp model, and the grey line corresponds to the line of linearity.
