## Appendix B: Supplementary Analysis for "Wearables-Only Analysis of Muscle and Joint Mechanics: An EMG-Driven Approach"

As discussed in the main text, an advantage of methods utilizing physics-based simulation (as in the proposed technique) over machine learning approaches is that clinically relevant internal state variables are necessarily characterized and the relationships between variables are dynamically consistent (i.e., they do not violate physical law). This was demonstrated in the main text by analyzing joint moment and muscle work. In this supplementary analysis, we compare estimates of other variables characterizing the muscle, MTU, and joint dynamics.

*Joint Kinematics*

Knee flexion angle was computed using an RTS Kalman smoother implementation of a previously validated complementary filter [1]. In deploying the proposed technique in practice, all data from a stride being analyzed would be available for analysis in which case optimal smoothing is preferred over filtering. Ankle dorsiflexion angle was estimated given the shank and foot orientation (as described in the main text) and following an Euler angle decomposition of the relative segment orientation. Estimation of the knee flexion angle ($r$ = 0.98, 4.08º RMSE) was better than for the ankle dorsiflexion angle ($r$ = 0.53, 9.93º). See Fig. B.1. for a graphical comparison.

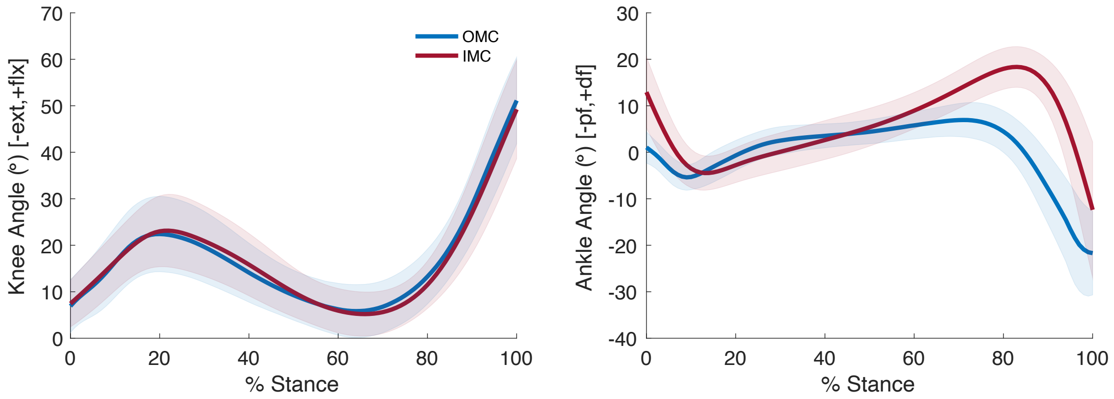


Fig. B.1. Ensemble average (± 1 s.d.) knee flexion angle (left) and ankle dorsiflexion angle (right) from OMC (blue) and IMC (red) analyses.

*MTU kinematics*

Figure B.2. and Table B.1. present the results of a comparison of MTU length estimates (computed as described in the main text and normalized by the length in the reference configuration) between OMC and IMC.

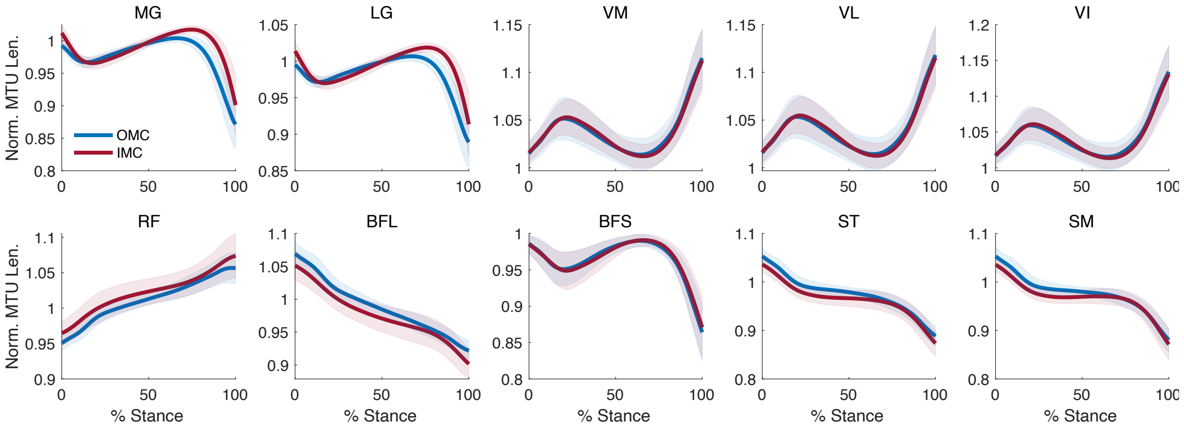


Fig. B.2. Ensemble average (± 1 s.d.) normalized MTU length from OMC (blue) and IMC (red) analyses.

Table B.1. Comparison of Norm. MTU Length: OMC vs IMC

| Muscle | $r$ | RMSE |
|---|---|---|
| MG | 0.83 | 0.02 |
| LG | 0.81 | 0.02 |
| VM | 0.98 | 0.01 |
| VL | 0.98 | 0.01 |
| VI | 0.98 | 0.01 |
| RF | 0.99 | 0.01 |
| BFL | 0.99 | 0.02 |
| BFS | 0.98 | 0.01 |
| ST | 0.99 | 0.02 |
| SM | 0.98 | 0.02 |

*Joint Power*

Net knee flexion power was computed as the product of knee joint velocity (flexion DOF, $\dot{\theta}$) and net knee flexion moment. Net knee flexion moment was computed as described in the main text and $\dot{\theta}$ was approximated numerically using the 5-point central difference method. Estimates from the proposed technique were moderately correlated ($r$ = 0.62) with those from ground truth inverse dynamics analysis with 0.53 W/kg (21.34 %range) RMSE and were strongly correlated ($r$ = 0.88) with those from OMC-Full with 0.37 W/kg (12.94 %range) RMSE (Fig. B.3.).

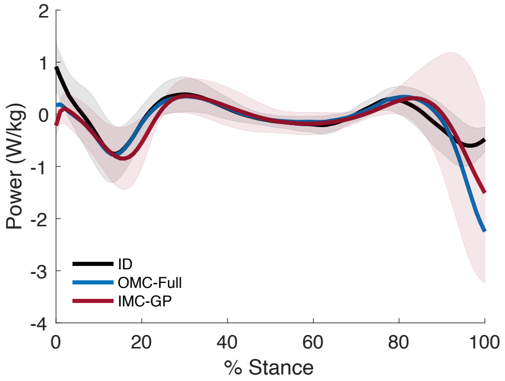

Fig. B.3. Ensemble average (± 1 s.d. shown for ID and IMC-GP) net knee flexion power from all three techniques.

*Muscle Power*

Figure B.4. and Table B.2. present the results of a comparison of muscle power (not to be confused with MTU power) between OMC-Full and IMC-GP computed as the product of the muscle force $f_m$ and the velocity $\dot{s}$ (see main text for definition of $s$).

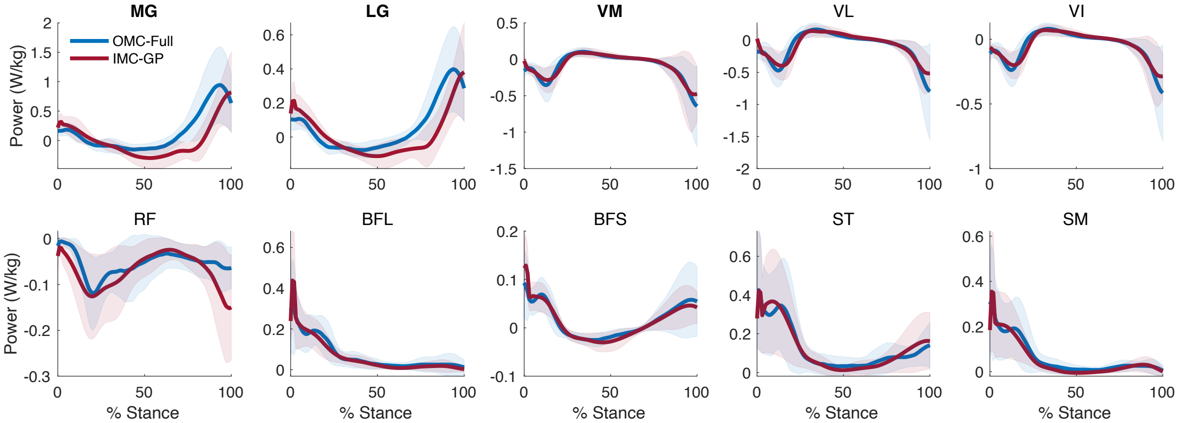

Fig. B.4. Ensemble average (± 1 s.d.) muscle power from OMC-Full (blue) and IMC-GP (red) analyses.

Table B.2. Comparison of Muscle Power: OMC-Full vs IMC-GP

| Muscle | *r* | RMSE (W/kg) |
|---|---|---|
| MG | 0.83 | 0.28 |
| LG | 0.85 | 0.11 |
| VM | 0.92 | 0.09 |
| VL | 0.88 | 0.16 |
| VI | 0.91 | 0.07 |
| RF | 0.73 | 0.04 |
| BFL | 0.89 | 0.05 |
| BFS | 0.91 | 0.02 |
| ST | 0.88 | 0.08 |
| SM | 0.90 | 0.05 |

*Muscle Force*

Figure B.5. and Table B.3. present the results of a comparison of muscle force between OMC-Full and IMC-GP computed as described in the main text.

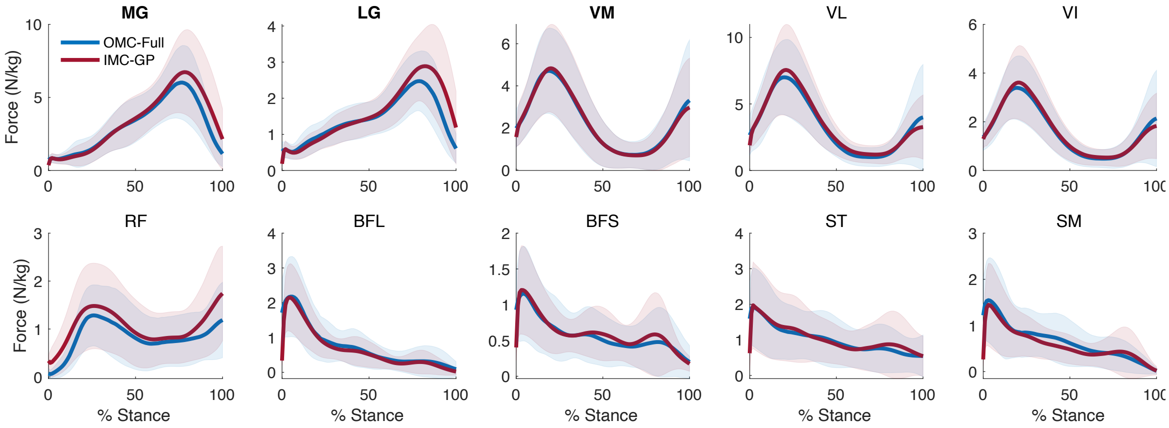


Fig. B.5. Ensemble average (± 1 s.d.) muscle force from OMC-Full (blue) and IMC-GP (red) analyses.

Table B.3. Comparison of Muscle Force: OMC-Full vs IMC-GP

| Muscle | *r* | RMSE (N/kg) |
|---|---|---|
| MG | 0.98 | 0.65 |
| LG | 0.97 | 0.36 |
| VM | 0.99 | 0.26 |
| VL | 0.96 | 0.87 |
| VI | 0.98 | 0.30 |
| RF | 0.86 | 0.41 |
| BFL | 0.90 | 0.38 |
| BFS | 0.80 | 0.27 |
| ST | 0.81 | 0.42 |
| SM | 0.85 | 0.32 |

*Muscle Moment*

Figure B.6. presents a graphical comparison of individual muscle moment between OMC-Full and IMC-GP. A statistical comparison was provided in the main text.

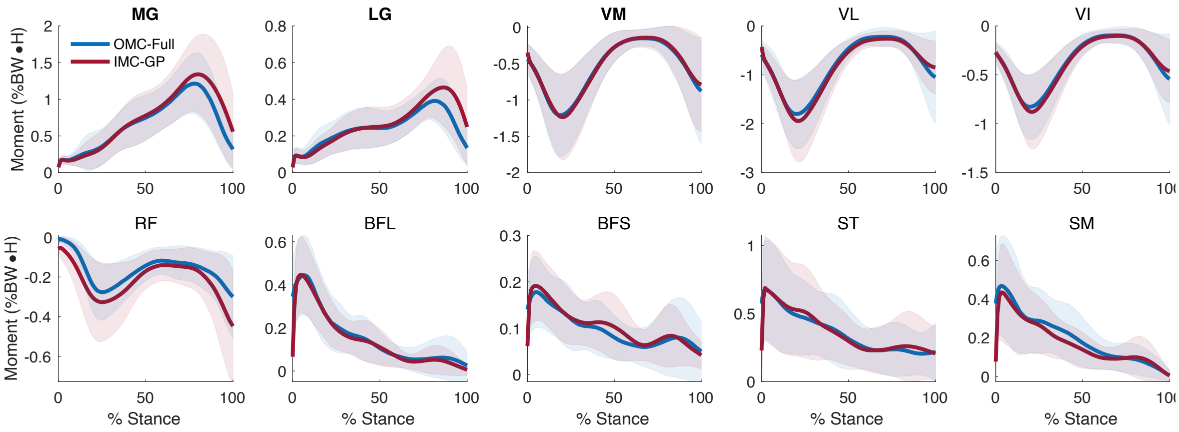


Fig. B.6. Ensemble average (± 1 s.d.) muscle moment from OMC-Full (blue) and IMC-GP (red) analyses.
